# Action at a distance: Defects in division plane positioning in the root meristematic zone affect cell organization in the differentiation zone

**DOI:** 10.1101/2021.04.30.442137

**Authors:** Alison M. Mills, Carolyn G Rasmussen

**Author notes:** Correspondence to Carolyn Rasmussen.

## Abstract

Cell division plane orientation is critical for plant and animal development and growth. TANGLED1 (TAN1) and AUXIN-INDUCED-IN-ROOT-CULTURES9 (AIR9) are division-site localized microtubule-binding proteins required for division plane positioning. *tan1* and *air9 Arabidopsis thaliana* single mutants have minor or no noticeable phenotypes but the *tan1 air9* double mutant has synthetic phenotypes including stunted growth, misoriented divisions, and aberrant cell-file rotation in the root differentiation zone. These data suggest that TAN1 plays a role in nondividing cells. To determine whether TAN1 is required in elongating and differentiating cells in the *tan1 air9* double mutant, we limited its expression to actively dividing cells using the G2/M-specific promoter of the syntaxin *KNOLLE* (*pKN:TAN1-YFP*). Unexpectedly, in addition to rescuing division plane defects, *pKN:TAN1-YFP* rescued root growth and the root differentiation zone cell file rotation defects in the *tan1 air9* double mutant. This suggests that defects that occur in the meristematic zone later affect the organization of elongating and differentiating cells.

**Summary Statement:** Expression of *TAN1* in the root meristematic zone rescues cell file rotation defects in *tan1 air9* mutants, suggesting defects that occur in mitosis may influence organization of nondividing cells.

## Introduction

Correct division plane orientation is key for patterning and growth across kingdoms. Because plant cells are confined by cell walls, division positioning is tightly regulated (Facette et al., 2018; Livanos and Müller, 2019; Rasmussen and Bellinger, 2018; Wu et al., 2018). Division plane determination begins during S or G2, when the nucleus is repositioned within the cell (Facette et al., 2018; Frey et al., 2010; Wada, 2018; Yi and Goshima, 2020). Polarity is often established and maintained by nuclear repositioning and polar localization of proteins during asymmetric division (Facette et al., 2018; Guo et al., 2021; Kimata et al., 2016; Muroyama and Bergmann, 2019; Shao and Dong, 2016; Wada, 2018). Next, land-plant cells typically form a structure around the nucleus at the cell cortex called the preprophase band (PPB). The PPB is a ring of microtubules, microfilaments and associated proteins that marks the future position of the new cell wall, called the division site (Li et al., 2015; Pickett-Heaps et al., 1999; Rasmussen and Bellinger, 2018; Smertenko et al., 2017; Van Damme, 2009). Nuclear and PPB positioning often match division predictions based on cell geometry (Martinez et al., 2018; Moukhtar et al., 2019). PPB disassembly upon nuclear envelope breakdown precedes spindle formation (Dixit and Cyr, 2002). After chromosome separation, the phragmoplast forms from the anaphase spindle to direct new cell wall synthesis. The phragmoplast is an antiparallel array of microtubules with plus-ends facing the cell center (Ho et al., 2012; McMichael and Bednarek, 2013; Müller and Jürgens, 2016). Kinesins transport vesicles to form the cell plate (Lee and Liu, 2013; Smertenko et al., 2018). New microtubule nucleation expands the phragmoplast outwards until the cell plate contacts the division site (Gu and Rasmussen, 2022; Murata et al., 2013; van Oostende-Triplet et al., 2017).

Division-site localized proteins including TANGLED1 (TAN1), PHRAGMOPLAST ORIENTING KINESIN1 (POK1), POK2, MICROTUBULE-ASSOCIATED PROTEIN 65-4 (MAP65-4), RAN GTPASE ACTIVATING PROTEIN (RAN-GAP), MYOSIN VIII and KINESIN-LIKE CALMODULIN BINDING PROTEIN (KCBP) remain at the cell cortex at the division site throughout cell division (Buschmann et al., 2015; Herrmann et al., 2018; Li et al., 2017; Lipka et al., 2014; Morgan et al., 2008; Walker et al., 2007; Wu and Bezanilla, 2014). Many of these proteins are important for division plane positioning, often during telophase. TAN1 is a division-site-localized protein required for phragmoplast guidance to the division site in maize (Martinez et al., 2017; Smith et al., 2001; Walker et al., 2007). TAN1 organizes microtubules at the cell cortex called cortical telophase microtubules which are incorporated into the phragmoplast to direct its movement at the cell cortex (Bellinger et al.). TAN1 binds and bundles microtubules in vitro (Martinez et al., 2020). Although the *tan1* maize mutant has misplaced divisions and stunted growth, *tan1 Arabidopsis thaliana* (Arabidopsis) mutants grow as well as wild-type plants and have minor division placement defects (Walker et al., 2007). Another division-site-localized protein, AIR9, also binds microtubules. AIR9 localizes to interphase cortical microtubule arrays, as well as co-localizing with the PPB, the phragmoplast, and localizing to the division site during late telophase (Buschmann et al., 2006). Similar to *tan1* single mutants, *air9* single mutants resemble wild-type plants (Buschmann et al., 2015). Due to their similar division-site localization, *tan1 air9* double mutants were generated in Arabidopsis. Combining mutations in both *tan1* and *air9* results in division-plane-positioning defects, stunted growth, and root twisting in the differentiation zone (Mir et al., 2018). While PPBs and phragmoplasts were both frequently misoriented in *tan1 air9* mutants, improper phragmoplast guidance is the primary defect (Mir et al. 2018). Transforming the *tan1 air9* double mutant with *TAN1-YFP* driven by the constitutive viral Cauliflower mosaic *CaMV35S* promoter rescues root growth, misoriented divisions, and cell-file-rotation defects (Mir et al., 2018).

We hypothesized that TAN1 may also have a role in organizing interphase microtubules in elongating and differentiated cells, because *tan1 air9* mutants had aberrant cell-file rotation in the root differentiation zone, minor defects in interphase microtubule organization, and root growth defects that were enhanced by the microtubule-depolymerizing-drug propyzamide (Mir et al., 2018). Cell file rotation phenotypes are often caused by mutations in microtubule-associated proteins or tubulin that alter the organization or stability of the interphase cortical microtubule array (Abe et al., 2004; Buschmann and Borchers, 2020; Buschmann et al., 2004; Hashimoto, 2015; Ishida et al., 2007; Nakajima et al., 2004; Sakai et al., 2008; Sedbrook et al., 2004; Shoji et al., 2004). For example, in several *alpha-tubulin* mutants, cell file rotation occurred in hypocotyls and root differentiation zones and in isolated cultured mutant cells (Abe et al., 2004; Buschmann et al., 2009; Ishida et al., 2007; Thitamadee et al., 2002). Cell file twisting also occurs when cell elongation differs between epidermal and cortical cells. Arabidopsis treated with compounds that affect microtubule stability, such as oryzalin or propyzamide, have helical cell files due to cortical cell swelling and reduced longitudinal cell expansion (Furutani et al., 2000; Hashimoto, 2002).

Therefore, defects in organ twisting are sometimes due to interphase microtubule disruption and likely independent of division-plane defects. However, several examples suggest that division-plane-orientation defects may lead to cell-file-rotation defects (Cnops et al., 2000; Wasteneys and Collings, 2009). Double mutants in two related receptor-like kinases have defects in division plane orientation near the quiescent center and in the endodermis and also have abnormal root skewing (Goff and Van Norman, 2021). Therefore, it is possible that either mitotic or non-mitotic defects lead to aberrant growth and root twisting defects.

To determine whether mitotic *TAN1* expression was sufficient to rescue root twisting in the differentiation zone of *tan1 air9* double mutants, we drove *TAN1* expression using the G2/M-phase-specific *KNOLLE* promoter (Lukowitz et al., 1996; Menges et al., 2005). KNOLLE is a syntaxin/Qa-SNARE required for cell-plate vesicle fusion (Strompen et al., 2002; Völker et al., 2001). The *KNOLLE* promoter drove *TAN1* expression in mitotic cells which rescued root growth and cell-file-rotation defects in the *tan1 air9* double mutant. Our results suggest that cell-file-rotation defects in the *tan1 air9* double mutant are likely due to defects that occur in actively dividing meristematic cells, and not due to a lack of TAN1 in nondividing cells.

## Results & Discussion

We generated two independent native-promoter TAN1 fluorescent protein fusions to determine whether *TAN1-YFP or CFP-TAN1* expressed by its native promoter would rescue the *tan1 air9* double mutant. Both constructs rescued the *tan1 air9* mutant. Previous studies showed that *35S*-driven *TAN1* expression rescued *tan1 air9* mutants (Mir et al., 2018). We drove expression of *CFP-TAN1* and *TAN1-YFP* using 1263 bp upstream of the start codon, *pTAN:CFP-TAN1* and *pTAN:TAN1-YFP*, and transformed or crossed them into the *tan1 air9* double mutant. Cell shape in the root tip (Figure 1A) and cell file rotation in the differentiation zone of *pTAN:CFP-TAN1 tan1 air9* plants was restored to *air9* single mutant levels (Figure 1B, 1C). Single *air9* mutants are indistinguishable from wild-type plants (Buschmann et al., 2015; Mir et al., 2018). Root cell division primarily occurs at the root tip (the meristematic zone). Above that, non-dividing cells elongate in the elongation zone. Root hairs mark the differentiation zone, where root cells mature and differentiate into different cell types (Wachsman et al. 2015). *tan1 air9* mutant roots tend to twist left with variable transverse cell-wall angle values that skew above 90° (Mir et al., 2018). *pTAN:CFP-TAN1* rescued *tan1 air9* root growth, with *pTAN:CFP-TAN1* expressing plants growing slightly longer than *air9* single mutants (Figure 1D). *pTAN:TAN1-YFP* also fully rescued *tan1 air9* root growth and restored normal root tip patterning (Supplementary Figure 1). Measuring PPB and phragmoplast angles is a metric for division plane orientation. PPB and phragmoplast angles were measured relative to the left-hand cell wall. *pTAN:CFP-TAN1* fully rescued PPB and phragmoplast positioning defects in *tan1 air9* mutants, restoring angle variances close to 90° (Figure 1E). This shows that mitotic expression of *TAN1* by its native promoter and fluorescent protein fusion at either end of the TAN1 protein is sufficient for normal plant growth, including the expansion and patterning of nondividing cells in the *tan1 air9* double mutant.

**Figure 1:**
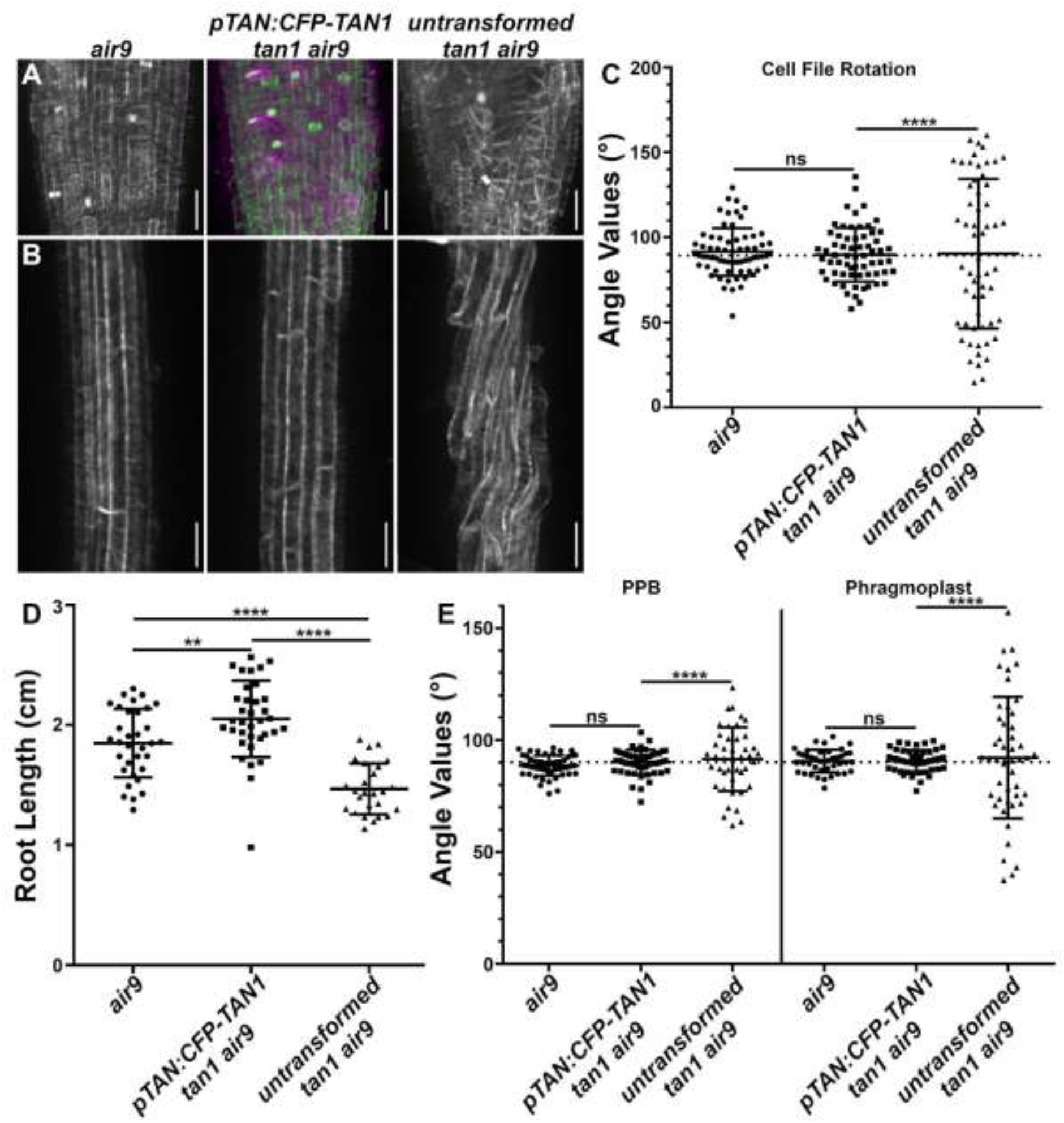
The TAN1 native promoter fused to TAN1 (*pTAN:CFP-TAN1*) rescues the *tan1 air9* double mutant. A) Maximum projections of 20 1-μm Z-stacks of root tips of an *air9, pTAN:CFP-TAN1* (magenta) *tan1 air9*, and untransformed *tan1 air9* plants expressing microtubule marker *UBQ10:mScarlet-MAP4* (green in the top middle panel). Bars = 25 μm. B) Maximum projections of 15 1-μm Z-stacks of the differentiation zone of an *air9*, *pTAN:CFP-TAN1 tan1 air9*, and untransformed *tan1 air9* plants expressing *UBQ10:mScarlet-MAP4*. Bars = 50 μm. C) Cell-file-rotation angles of *air9, pTAN:CFP-TAN1 tan1 air9*, and untransformed *tan1 air9* plants, n>11 plants for each genotype and n>57 cells for angle measurements. Cell-file-angle variances were compared with Levene’s test due to the non-normal distribution. D) Root-length measurements from 8 days after stratification of *air9, pTAN:CFP-TAN1 tan1 air9*, and untransformed *tan1 air9* plants, n>25 plants for each genotype, compared by two-tailed t-test with Welch’s corrections. E) PPB and phragmoplast angle measurements in *air9, pTAN:CFP-TAN1 tan1 air9*, and untransformed *tan1 air9* plants, n>9 plants for each genotype. N>41 cells for angle measurements. PPB and phragmoplast angle variations compared with F-test. ns indicates not significant, ** P-value <0.01, **** P-value <0.0001.

Previous fluorescence measurements of TAN1-YFP in wild-type lines expressing *pTAN:TAN1-YFP* demonstrated that fluorescent signal above background was limited to the meristematic zone (Mir et al., 2018). We hypothesized that TAN1 accumulated at low but undetectable levels in interphase cells when driven by its native promoter. To test whether *TAN1* expression limited to mitotic cells influenced root growth and suppressed root twisting in the *tan1 air9* double mutant, we fused the *KNOLLE* promoter to *TAN1-YFP* (*pKN:TAN1-YFP*) and transformed it into the *tan1 air9* double mutant. The *KNOLLE* promoter is specifically expressed in G2/M and is contingent on the MYB (myeloblastosis) transcription factors MYB3R1 and MYB3R4 which promote mitosis-specific gene expression (Haga et al., 2011; Yang et al., 2021). Our prediction was that *pKN:TAN1-YFP* would fully rescue mitotic defects but not restore root growth or suppress aberrant cell file rotation within the root differentiation zone in the *tan1 air9* mutant.

*pKN:TAN1-YFP* fully rescued the defects in *tan1 air9* mutants (Figure 2, other independent lines in Supplementary Figure 2). This includes rescuing cell patterning and cell-file-rotation defects (Figure 2A-C), root growth (Figure 2D), and PPB and phragmoplast positioning (Figure 2E). In addition, *pKN*-drivenTAN1-YFP localized to the division site during mitotic stages similar to *pTAN1*-driven CFP-TAN1 (Supplementary Figure 3). We compared phenotypes of *pKN:TAN1-YFP* to the *35S:TAN1-YFP* lines which rescue the *tan1 air9* mutant (Mir et al. 2018). Both *35S:TAN1-YFP* and *pKN:TAN1-YFP* significantly rescued the *tan1 air9* double mutant (Figure 3A & 3B). Root growth and PPB and phragmoplast angles were equivalent in *tan1 air9* plants expressing *pKN:TAN1-YFP* or *35S:TAN1-YFP* (Figure 3D & 3E). However, *pKN:TAN1-YFP* reduced cell-file-rotation variability slightly more than *35S:TAN1-YFP* (Figure 3C). This suggests that expressing *TAN1* in dividing cells is sufficient to fully rescue the *tan1 air9* double mutant. To determine why rescue with the *KNOLLE* promoter resulted in less cell-file-rotation variance, we measured TAN1-YFP fluorescence intensities in the *35S:TAN1-YFP* and *pKN:TAN1-YFP* lines. *pKN:TAN1-YFP* was expressed strongly in the meristematic zone of root tips (Figure 4B), often showing TAN1-YFP fluorescence in recently divided cells, similar to native promoter driven accumulation (Figure 1, Supplementary Figure 1, (Mir et al., 2018)). Indeed, TAN1-YFP accumulated at higher levels in the meristematic zone when expression was driven by the *KNOLLE* promoter (Figure 4G). However, unlike TAN1-YFP from *p35S:TAN1-YFP*, TAN1-YFP did not accumulate above background levels in the elongation and differentiation zone of roots expressing *pKN:TAN1-YFP* (Figure 4F & 4G). Lack of TAN1-YFP outside the meristematic zone and more complete rescue of *tan1 air9* cell file rotation by *pKN:TAN1-YFP* suggests that TAN1 is not required in elongating and differentiating cells. In other words, cell-file-rotation defects may be a consequence of defects that occur within the root meristematic zone either during mitosis or shortly afterwards.

**Figure 2:**
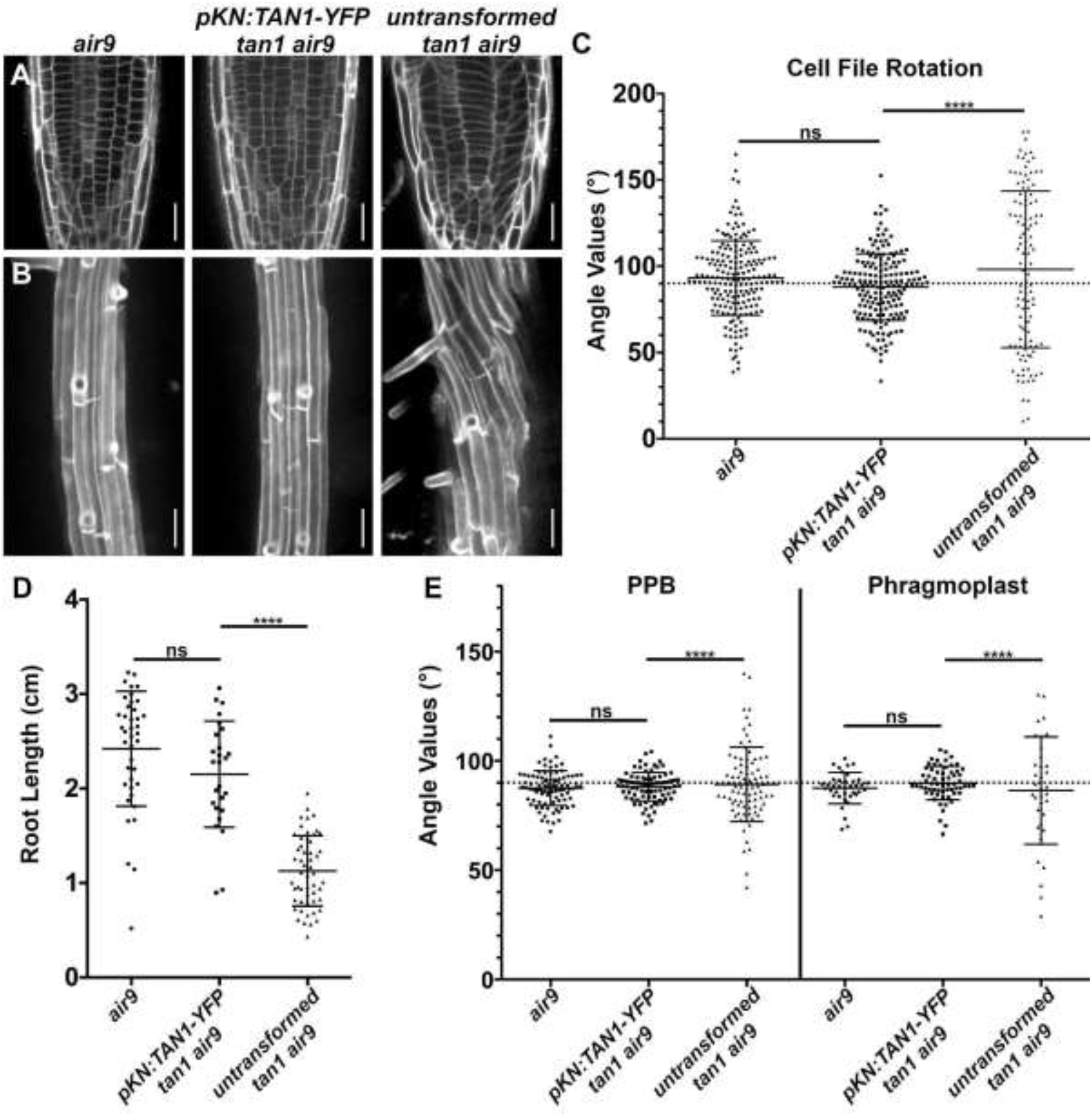
Full rescue of the *tan1 air9* double mutant with the G2/M-specific *KNOLLE* promoter fused to TAN1 (*pKN:TAN1-YFP*). A) Propidium iodide (PI) stained cell walls in root tips of an *air9, pKN:TAN1-YFP tan1 air9*, and untransformed *tan1 air9* plants. Bars = 25 μm. B) Maximum projections of 10 1-μm Z-stacks of PI-stained differentiation zone root cell walls. Bars = 50 μm. C) Cell-file-rotation angles of *air9, pKN:TAN1-YFP tan1 air9*, and untransformed *tan1 air9* plants, n>23 plants for each genotype. N>114 cells for angle measurements. Cell-file-rotation angle variances were compared with Levene’s test due to the non-normal distribution. D) Root-length measurements from 8 days after stratification of *air9, pKN:TAN1-YFP tan1 air9*, and untransformed *tan1 air9* plants, n>25 plants for each genotype, compared by two-tailed t-test with Welch’s corrections. E) PPB and phragmoplast angle measurements in *air9, pKN:TAN1-YFP tan1 air9*, and untransformed *tan1 air9* plants, n>20 plants for each genotype. N>34 cells for angle measurements. PPB and phragmoplast angle variations compared with F-test. ns indicates not significant, **** P-value <0.0001.

**Figure 3:**
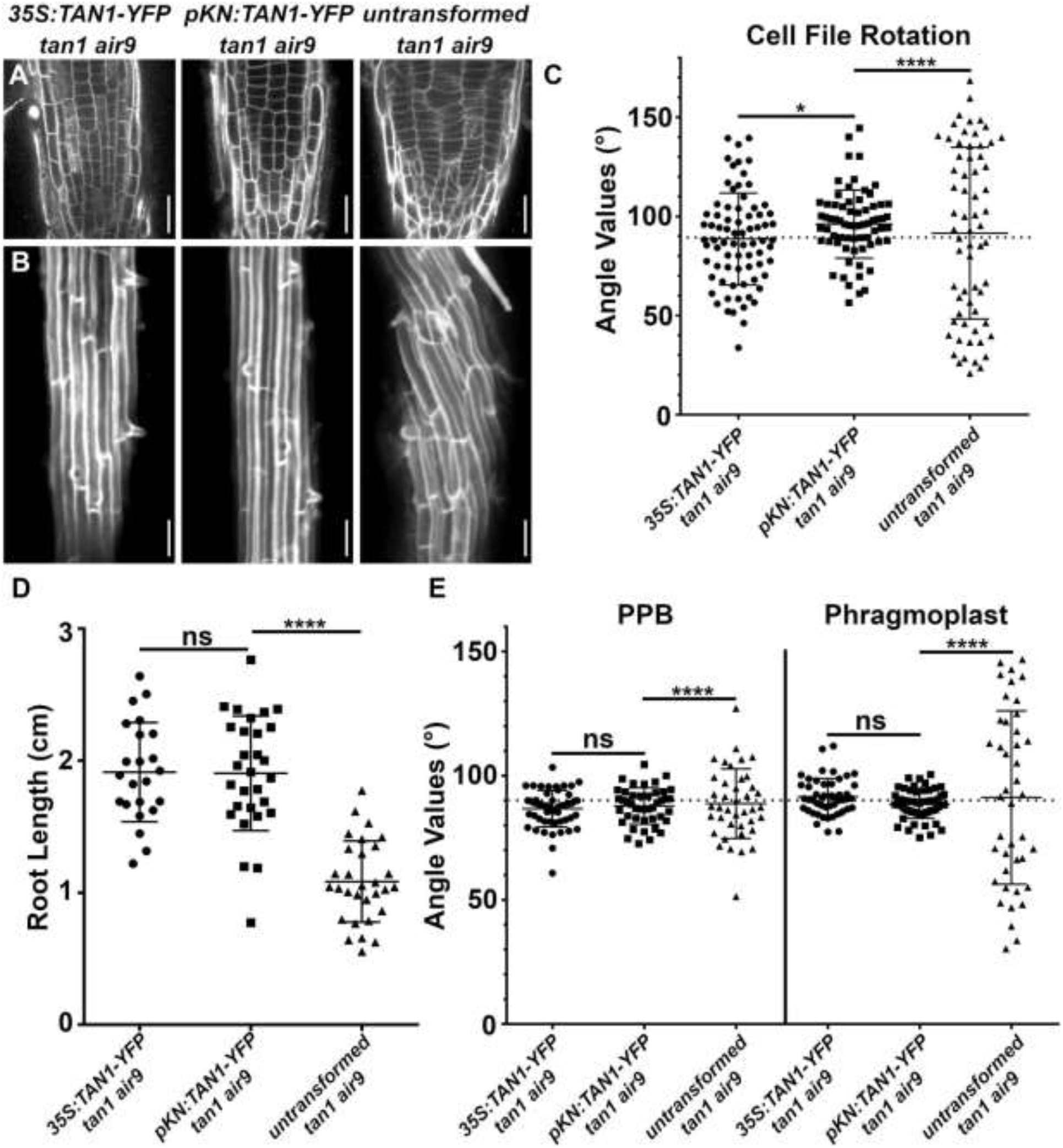
Comparison between *KNOLLE* promoter driven (*pKN:TAN1-YFP*) and *35S* driven TAN1 (*p35S:TAN1-YFP*) rescue of the *tan1 air9* double mutants. A) Propidium iodide (PI) stained root tips of *tan1 air9* mutants expressing *p35S:TAN1-YFP, pKN:TAN1-YFP*, and untransformed plants. Bars = 25 μm. B) Maximum projections of 10 1-μm Z-stacks of PI-stained differentiation-zone root cell walls of *tan1 air9* mutants expressing *p35S:TAN1-YFP, pKN:TAN1-YFP*, and untransformed plants. Bars = 50 μm. C) Cell-file-rotation angles of *tan1 air9* mutants expressing *p35S:TAN1-YFP, pKN:TAN1-YFP*, and untransformed plants, n>13 plants for each genotype. N>64 cells for angle measurements. Angle variances were compared with Levene’s test. D) Root-length measurements from 8 days after stratification of *tan1 air9* mutants expressing *p35S:TAN1-YFP, pKN:TAN1-YFP*, and untransformed plants, n>17 plants for each genotype, compared by two-tailed t-test with Welch’s corrections. E) PPB and phragmoplast angle measurements in *tan1 air9* double mutants expressing *p35S:TAN1-YFP, pKN:TAN1-YFP*, and untransformed plants, n>12 plants for each genotype. N>39 cells for angle measurements. PPB and phragmoplast angle variations compared with F-test. ns indicates not significant, * P-value <0.05, **** P-value <0.0001.

**Figure 4:**
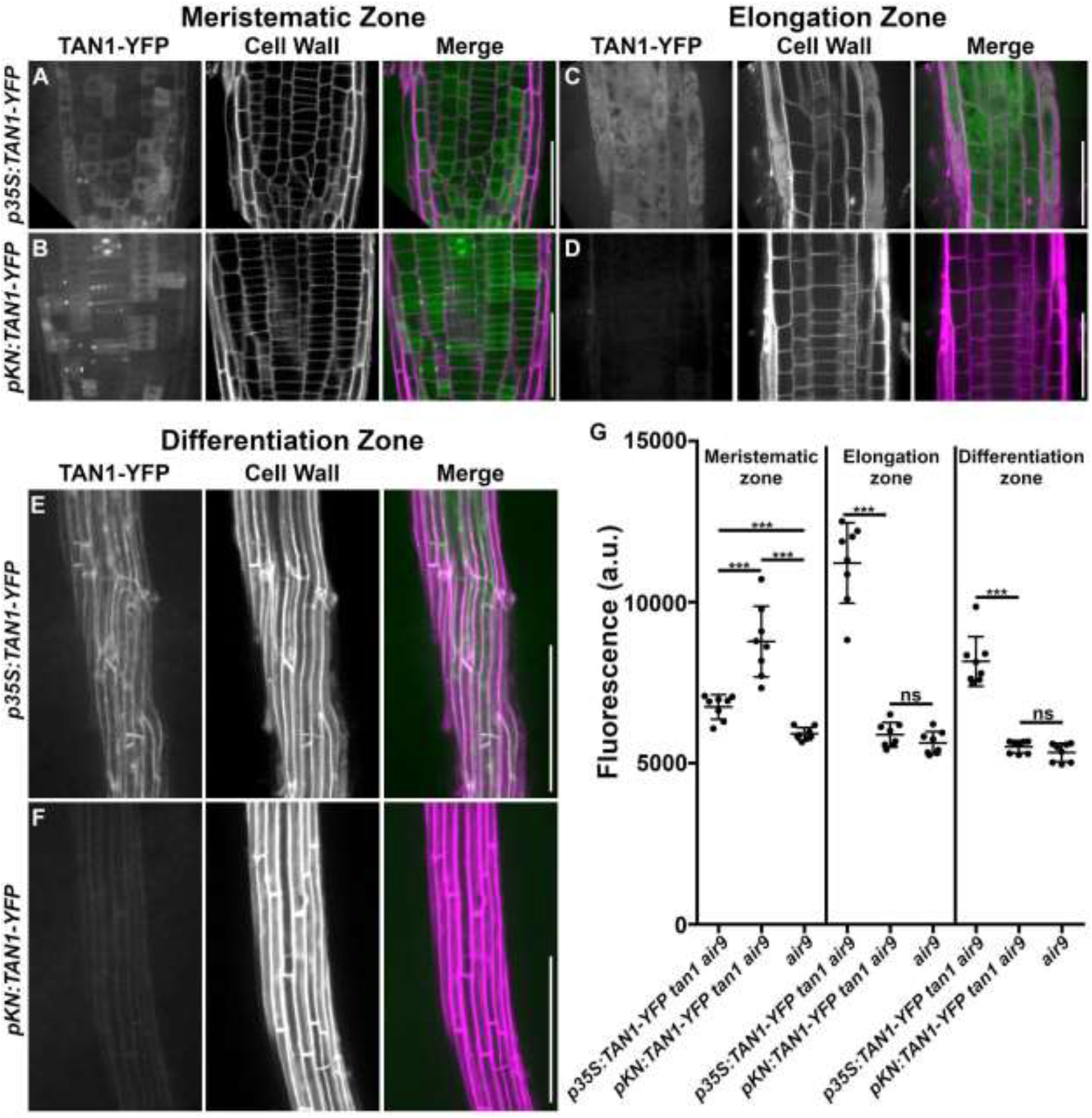
Comparison of TAN1-YFP fluorescence intensity when driven by the constitutive *35S* promoter (*p35S:TAN1-YFP*) and G2/M-specific *KNOLLE* promoter (*pKN:TAN1-YFP*) in *tan1 air9* roots. (A and B) Micrographs of the meristematic zone, (C and D) maximum projections of 3 1-μm Z-stacks of the elongation zone, and (E and F) maximum projections of 10 1-μm Z-stacks of the differentiation zone of *tan1 air9* mutants expressing (A, C, E) *p35S:TAN1-YFP* or (B, D, F) *pKN:TAN1-YFP*. Cell walls were stained with propidium iodide. Root tip and elongation zone, bars = 50μm. Differentiation zone, bars = 200μm. G) TAN1-YFP fluorescence-intensity measurements (arbitrary units, a.u.) from the meristematic zone, elongation zone, and differentiation zone of *tan1 air9* mutants expressing *p35S:TAN1-YFP, pKN:TAN1-YFP*, and *air9* mutants, n=8 plants for each genotype, fluorescence compared with Mann-Whitney U test. *** P-value <0.001. ns indicates not significant.

Another example of defects in mitotic expression and division plane positioning affecting nondividing cell organization occurs in the MYB activated GRAS-type (GIBBERELLIC-ACID INSENSITIVE, REPRESSOR of GAI and SCARECROW-type) transcription factor *scarecrow-like 28-3* (*scl28-3*) mutant. G2/M specific gene expression controlled by SCL28 is important for mitotic progression and division-plane positioning. *scl28-3* mutants have both misoriented divisions and root twisting (Goldy et al., 2021).

How do defects that occur within the meristematic zone influence the patterning or shape of nondividing, differentiating root cells and root growth? Our hypothesis is that misshapen cells and improper division plane orientation in *tan1 air9* double mutants cause the uneven distribution of mechanical stresses across the root, which then triggers cell wall integrity responses that limit growth and alter root organization. Cell wall stress patterns depend on cell geometry and the mechanical properties of cell walls (Cosgrove, 2018; Hamant and Haswell, 2017; Whitewoods and Coen, 2017; Schopfer, 2006).

Division-plane positioning is a way plants may respond to mechanical stress (Chakrabortty et al., 2018; Louveaux et al., 2016). Cell division relieves mechanical stress by creating smaller cells with less surface; further, divisions along maximal tensile stress promote growth homogeneity (Alim et al., 2012; Sapala et al., 2018). Microtubules often align parallel to maximal tensile stress (Hamant et al., 2008; Heisler et al., 2010; Sampathkumar et al., 2014; Uyttewaal et al., 2012) and cortical-microtubule alignment often influences PPB placement (Louveaux et al., 2016; Rasmussen et al., 2013; Wick and Duniec, 1983). However, division plane positioning is disrupted in mutants with division plane orientation defects. Although *tan1 air9* cells may perceive mechanical stress, phragmoplast guidance defects prevent construction of new cell walls in an orientation that minimizes mechanical stress. Abnormal stresses are perceived by receptor-like kinases involved in the cell-wall-integrity response. Cell-wall-integrity responses trigger slow growth, upregulation of stress responses, and changes in cell morphogenesis, (Buschmann and Borchers, 2020; Caño-Delgado et al., 2003; Gonneau et al., 2018; Hématy et al., 2007; Wolf et al., 2014), which may contribute to the stunted growth and twisted cell files observed in the *tan1 air9* double mutant.

## Materials and methods

### Plasmid Construction

*pKN:TAN1-YFP* was generated by amplifying 2152 bp of the 5’ *KNOLLE* (AT1G08560) promoter from Columbia with primers pKN-5’SacI Fw and pKN-5’EcoRI Rw. EcoRI and StuI double digestion was used to introduce the *KNOLLE* promoter into pEZT-NL containing the TAN1 coding sequence. Primers 35SpKN5’ Fw and YFP XhoI Rw were used to amplify *pKN:TAN1-YFP* then XhoI and StuI double digestion was used to clone *pKN:TAN1-YFP* into pEGAD, a gift from Professor Sean Cutler (University of California, Riverside).

*pTAN:CFP-TAN1* was created by overlapping PCR. The 1263bp 5’ sequence upstream of genomic *TAN1* was amplified using *Np:AtTAN-YFP* (Walker et al., 2007) as a template with the primers NpTANSacIFor and NpTANceruleanRev. Cerulean fluorescent protein (CFP) was amplified using Cerulean CDS in pDONR221P4r/P3r, a kind gift from Professor Anne Sylvester (University of Wyoming), as template with the primers NpTANceruleanFor and CeruleanpEarleyRev. TAN1 CDS was amplified using *35S:YFP-TAN1* in pEarley104 as a template with the primers CeruleanpEarleyFor and pEarleyOCSPstIRev. The 1263bp TAN1 native promoter, CFP, and TAN1 CDS were combined to create *pTAN:CFP-TAN1* by overlapping PCR with the primers NpTANSacI and pEarleyOCSPstIRev. SacI and PstI double digest was used to subclone *pTAN:CFP-TAN1* into pJHA212G, a kind gift from Professor Meng Chen (University of California, Riverside).

### Generation of Transgenic Lines

Transgenic Arabidopsis lines were generated using *Agrobacterium tumefaciens*-mediated floral dip transformation as described (Clough and Bent, 1999). Previously described *tan1 air9* mutants (Mir et al., 2018), *csh-tan* (*TAN1*, AT3G05330 (Walker et al., 2007)) and *air9-31* (*AIR9*, AT2G34680 (Buschmann et al., 2015)), were used for floral dip transformation of *pKN:TAN1-YFP* and T1 transgenic plants were subsequently selected on 1/2 MS plates containing 15 μg/mL glufosinate (Finale; Bayer). TAN1-YFP signal in T1 plants was confirmed by confocal microscopy before being transferred to soil and selfed. The genotypes of *csh-tan1 air9-31* transformants was confirmed using the primers AIR9_cDNA 2230 F and AIR9 gnm7511 R (to identify *AIR9* wild-type), AIR9 gnm7511 R and Ds5-4 (to identify T-DNA insertion in *AIR9*), ATLP and AtTAN 733-CDS Rw (to identify TAN1 wild-type), and AtTAN 733-CDS Rw and Ds5-4 (to identify T-DNA insertion in *TAN1*). The microtubule marker *CFP-TUBULIN* (Kirik et al., 2007), a kind gift from Professor David Erhardht (Stanford University) was crossed into *pKN:TAN1-YFP tan1 air9* plants using *tan1 air9 CFP-TUBULIN* plants (Mir et al., 2018).

*air9-5 tan-mad* Columbia/Wassilewskija double mutants (Mir et al., 2018) expressing the microtubule marker *UBQ10:mScarlet-MAP4* (Pan et al., 2020), *a* kind gift from Professor Zhenbiao Yang (University of California, Riverside), was used for floral dip transformation of *pTAN1:CFP-TAN1* and selected on 1/2 MS plates containing 100 μg/mL gentamicin (Fisher Scientific). T1 seedlings were screened for mScarlet and CFP signal and then transferred to soil to self.

### Growth conditions and root length measurements

Plates containing ½ strength Murashige and Skoog (MS) media (MP Biomedicals; Murashige and Skoog, 1962) containing 0.5 g/L MES (Fisher Scientific), pH 5.7, and solidified with 0.8% agar (Fisher Scientific) were used to grow Arabidopsis seedlings. *tan1 air9* transgenic lines expressing T3 *p35S:TAN1-YFP*, T2 *pKN:TAN1-YFP*, and T2 *pTAN:CFP-TAN1* were used for root length experiments. At least 3 biological replicates were used for each root growth assay. 5-7 1/2 MS plates were used for each replicate. 12-15 seeds were sown in a single level line on each plate with untransformed *tan1 air9* double mutants and *air9* single mutants sown on plates alongside double mutants expressing *TAN1* constructs. Seeds were stratified on plates in the dark at 4°C for 2 to 5 days. After stratifying, plates were positioned vertically in a growth chamber (Percival) with a 16/8-h light/dark cycle and temperature set to 22°C. Each biological replicate was placed in the growth chamber on different days. 8 days after stratification, plates were scanned (Epson) and root lengths were measured using FIJI (ImageJ, http://fiji.sc/). Transgenic seedlings were screened for fluorescence by confocal microscopy to identify seedlings expressing YFP, CFP and mScarlet translational fusion transgenes when present. Each root growth experiment had a minimum of 3 biological replicates. Statistical analysis of root length was determined using Welch’s t-test with Prism (GraphPad) and replicates were checked for discrepancies in statistical significance before pooling replicates for analysis. Root length plots were created using Prism (GraphPad).

To assess the ability of *TAN1* driven by its native promoter to rescue the *tan1 air9* double mutant, *Np:AtTAN-YFP* (Walker et al., 2007) was crossed to *tan-mad air9-5* double mutants. The progeny of *pTAN1:TAN1-YFP tan-mad*/+ *air9-5*/+ plants were sown on 1/2 MS media and grown as described above. The seedlings were screened by confocal microscopy for the presence of TAN1-YFP and then collected for genotyping. Seedlings were genotyped with primers AtExon1_1For and At255AfterStopRev (to identify TAN wild-type), JL202 and ATLP (to identify T-DNA insertion in *TAN1*), AIR9-5RP and AIR9-5LP (to identify wild-type *AIR9*), and LBb1.3 and AIR9RP (to identify T-DNA insertion in *AIR9*) (Supplementary Table 1). The length of *tan1 air9* double mutants expressing *pTAN1:TAN1-YFP* was compared to *tan1 air9* double mutants and *air9* single mutant siblings lacking *pTAN1:TAN1-YFP*. *air9* single mutants used for root length analysis included *air9/air9 TAN1/TAN1* and *air9/air9 TAN1/tan1* plants.

### Confocal Microscopy

Imaging and screening was performed using Micromanager software (micromanager.org) running an inverted Ti Eclipse (Nikon) with motorized stage (ASI Piezo) and spinning-disk confocal microscope (Yokogawa W1) built by Solamere Technology. Solid-state lasers (Obis) were used with standard emission filters (Chroma Technology). Excitation 445, emission 480/40 (for CFP-translational fusions); excitation 514, emission 540/30 (for YFP-translational fusions); and excitation 561, emission 620/60 (for propidium iodide and mScarlet-MAP4) were used. The 20x objective has a 0.75 numerical aperture. The 60x objective was used with perfluorocarbon immersion liquid (RIAAA-6788, Cargille) and has a 1.2 numerical aperture objective.

### Measurements of PPB and phragmoplast angles and cell file rotation

All angle data was gathered from at least 3 biological replicates. Each replicate consisted of 5-7 1/2 MS plates with 12-15 seeds sown on each plate. 4-5 seeds of each genotype were sown on each plate to ensure growing conditions were identical. Each replicate was moved from stratifying to the growth chamber on independent days. Seedlings were then imaged at 8 days after stratification. Images were collected by confocal microscopy using the 20x objective to collect images of the differentiation zone for cell file angles and the 60x objective to collect images of root tips expressing a microtubule marker (*CFP-TUBULIN* or *mScarlet-MAP4*) for PPB and phragmoplast angles. The differentiation zone was identified by the presence of root hairs. Angles were measured using FIJI. Cell file angles were measured from the left hand side of the cell taking the angle between the long axis of the root and the transverse cell wall using images of root cells in the differentiation zone. PPB and phragmoplast angles were measured by taking the angle between the left-hand cell wall and the PPB or phragmoplast. Because CFP-TUBULIN was faint and poorly marked cell boundaries, the cell walls of *CFP-TUBULIN* expressing seedlings were stained with 10 μM PI for 1 minute before destaining in distilled water prior to imaging. Each angle measurement represents a single angle measured from one cell.

Statistical analyses were performed using Excel (Microsoft Office) and Prism (GraphPad). To compare normally distributed variance of PPB and phragmoplast angles F-test was used. Levene’s test was used to compare variances of cell file angle measurements because *tan1 air9* cell file angles are non-normally distributed due to left hand twisting of the roots. Angle variance across biological replicates was checked before pooling data.

### Fluorescence Intensity Measurements

*air9, 35S:TAN1-YFP tan1 air9*, and *pKN:TAN1-YFP tan1 air9* plants were grown on 1/2 MS plates as described above. 8 days after stratification, plants were imaged by confocal microscopy using identical settings. Root tips were imaged using the 60X objective. The median fluorescence intensity of an 116,001.5 μm^2^ area was measured from multiple individual plants of each genotype. Each fluorescence measurement represents the median fluorescence from a single meristematic zone from one plant. Elongation zone and differentiation zone images were taken with the 20x objective and the median fluorescence intensity of an 12,323.4 μm^2^ area was measured from multiple individual plants of each genotype. Each fluorescence measurement represents the median fluorescence from a single elongation or differentiation zone from one plant.

## Acknowledgements

We thank Stephanie Martinez (University of California, Riverside, UCR) and Aimee Uyehara (UCR) for improving manuscript clarity, Professors Henrik Buschmann (Osnabruck University), Meng Chen (UCR), Zhenbio Yang (UCR), Sean Cutler (UCR), David Ehrhardt (Carnegie Institute), Anne Sylvester (University of Wyoming) and Dr. Ricardo Mir (UCR) for materials or facilities, and Professor Jaimie Van Norman (UCR) for cloning advice. Funding from NSF-MCB grant 1716972, NSF-CAREER grant 1942734, and USDA CA-R-BPS-5108-H is gratefully acknowledged.

## Supplementary Figures 1–3 and Supplementary Table

**Supplementary Figure 1.**
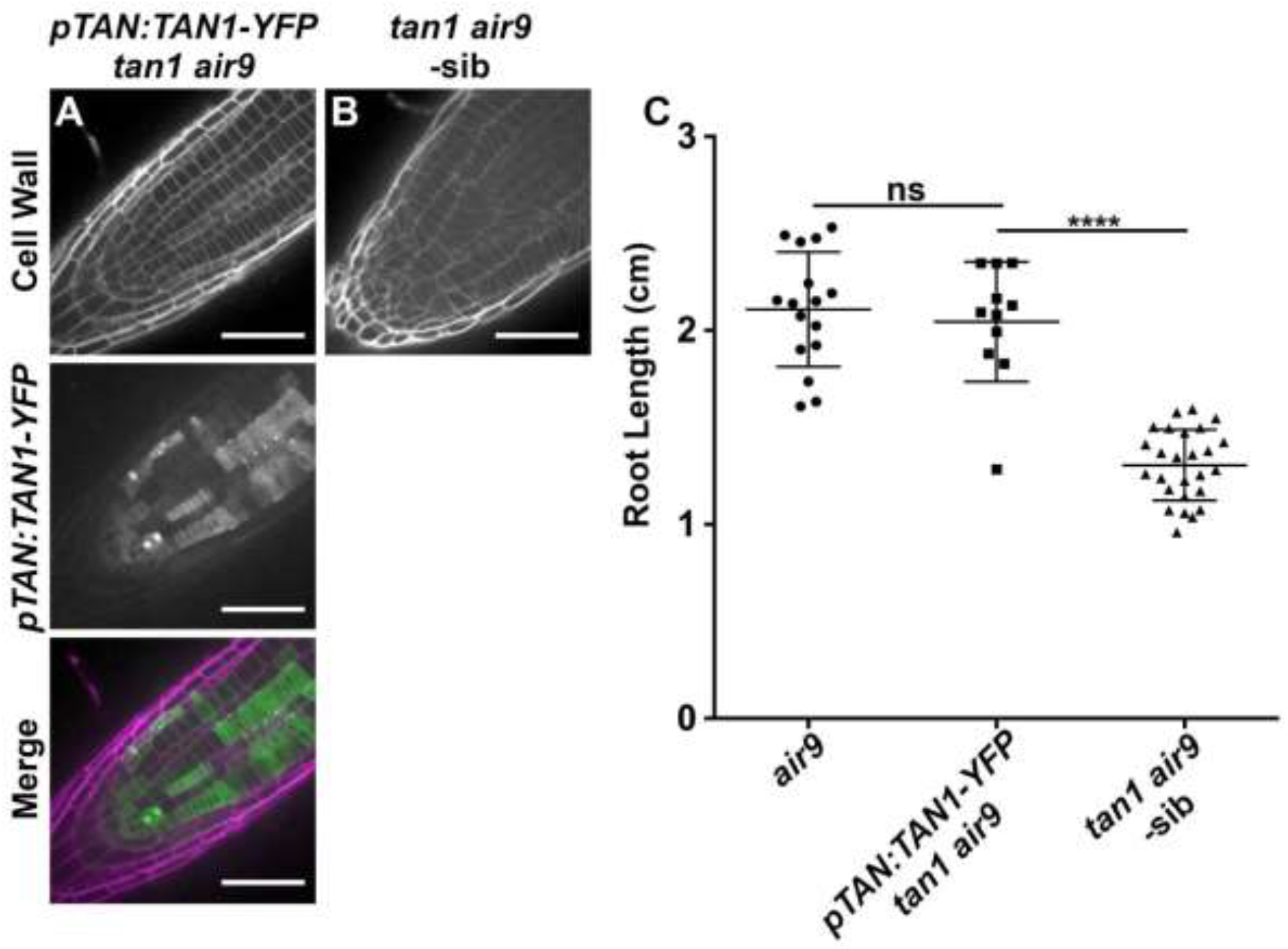
*TAN1-YFP* expressed by its native promoter (*pTAN:TAN1-YFP*) rescues *tan1 air9* double mutant root growth. Confocal images of propidium iodide-stained roots of *tan1 air9* plants. A) A *tan1 air9* plant expressing *pTAN:TAN1-YFP*. B) A negative sibling *tan1 air9* plant. Bars = 50 μm. C) Root length measurements from 8 days after stratification of *air9* single mutants (left), *pTAN1:TAN1-YFP tan1 air9* double mutants (middle), and *tan1 air9* double mutants (right). n > 10 plants for each genotype, compared by two-tailed t-test with Welch’s correction. ns indicates not significant, **** P-value <0.0001.

**Supplementary Figure 2.**
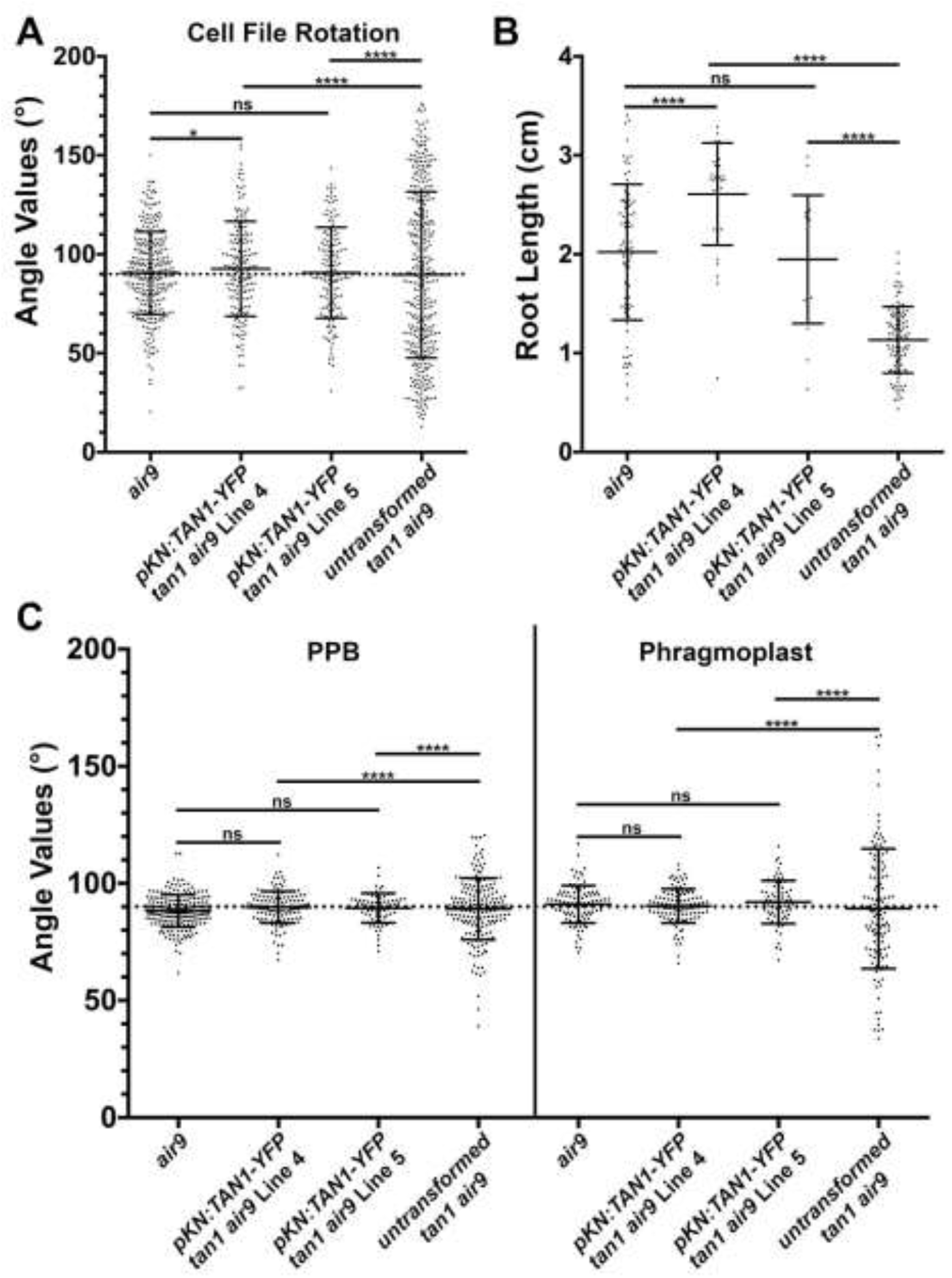
*pKN:TAN1-YFP tan1 air9* lines show significant rescue compared to untransformed *tan1 air9*. A) Cell file rotation angles of *air9* single mutants (left), two transgenic lines expressing pKN:*TAN1-YFP* in the *tan1 air9* double mutant designated as line 4 (center left) and line 5 (center right) and untransformed plants (right), n >17 plants for each genotype. N > 146 cells for angle measurements. Angle variances were compared with Levene’s test. B) Root length measurements from 8 days after stratification of *air9* single mutants (left), two transgenic lines expressing pKN:*TAN1-YFP* in the *tan1 air9* double mutant (middle), and untransformed plants (right), n > 21 plants for each genotype, compared by two-tailed t-test with Welch’s correction. C) PPB and phragmoplast angle measurements in dividing root cells of *air9* single mutants (left), two transgenic lines expressing pKN:*TAN1-YFP* in the *tan1 air9* double mutant (middle), and untransformed plants (right), n > 15 plants for each genotype. N > 69 cells for angle measurements. Angle variations compared with F-test. ns indicates not significant, * P-value < 0.05, **** P-value <0.0001.

**Supplementary Figure 3.**
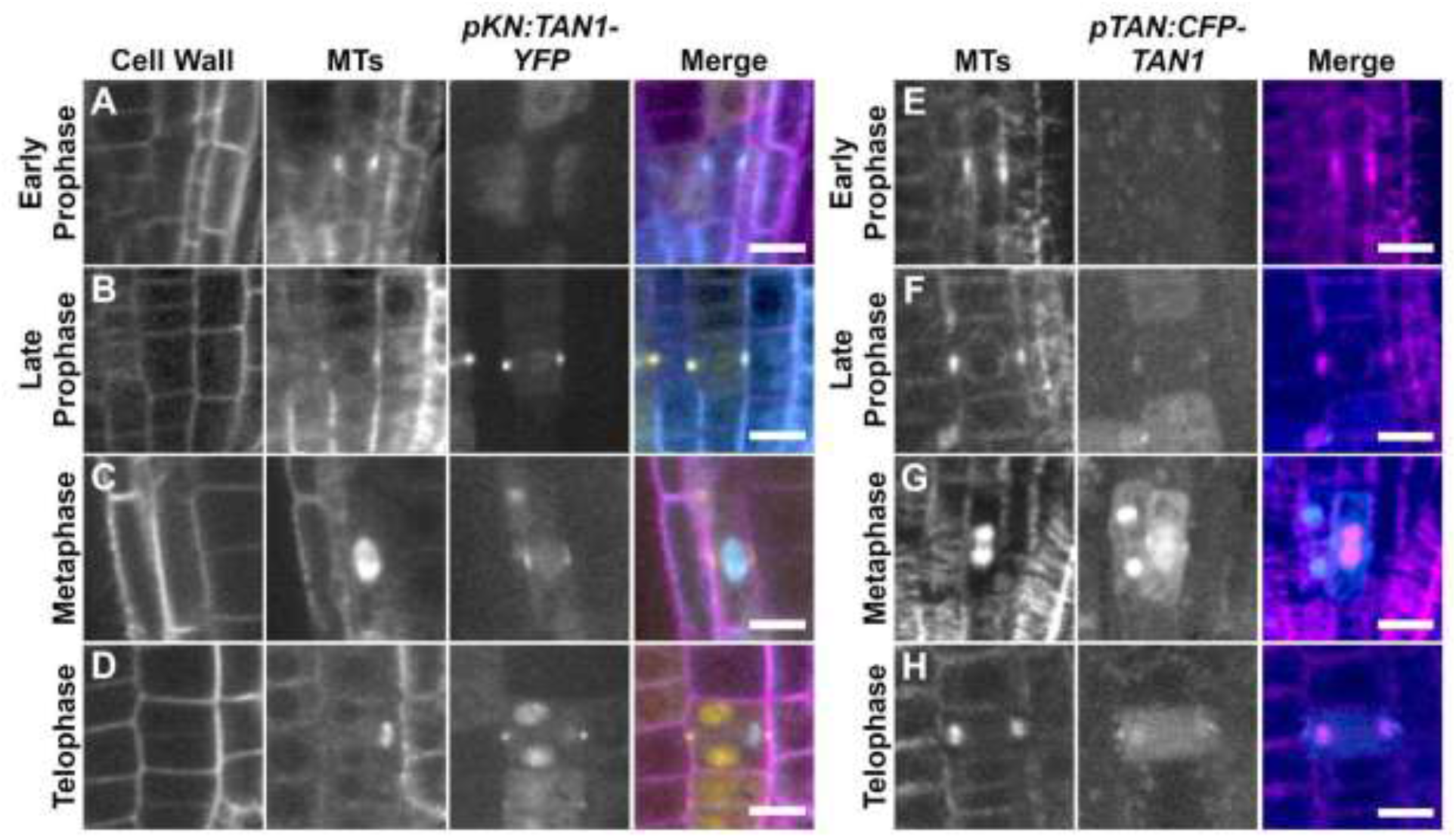
Division site localization of TAN1-YFP driven by the *KNOLLE* promoter (*pKN:TAN1-YFP*) and CFP-TAN1 driven by the *TAN1* promoter (*pTAN:CFP-TAN1*) in *tan1 air9* double mutants. A-D) Confocal images of propidium iodide-stained (Cell Wall) roots of *tan1 air9* plants expressing *pKN:TAN1-YFP* and *CFP-TUBULIN* (MTs) in dividing root tip cells. E-F) Maximum projections of 3 1-μm Z-stacks of *tan1 air9* plants expressing *pTAN:CFP-TAN1* and the microtubule (MTs) marker *UBQ10:mScarlet-MAP4* in dividing root tip cells. Representative images of cells with (A&E) broad early PPBs, (B&F) late narrow PPBs, (C&G) metaphase spindles, and (D&H) phragmoplasts. Bars = 10 μm.

**Supplementary Table 1.**
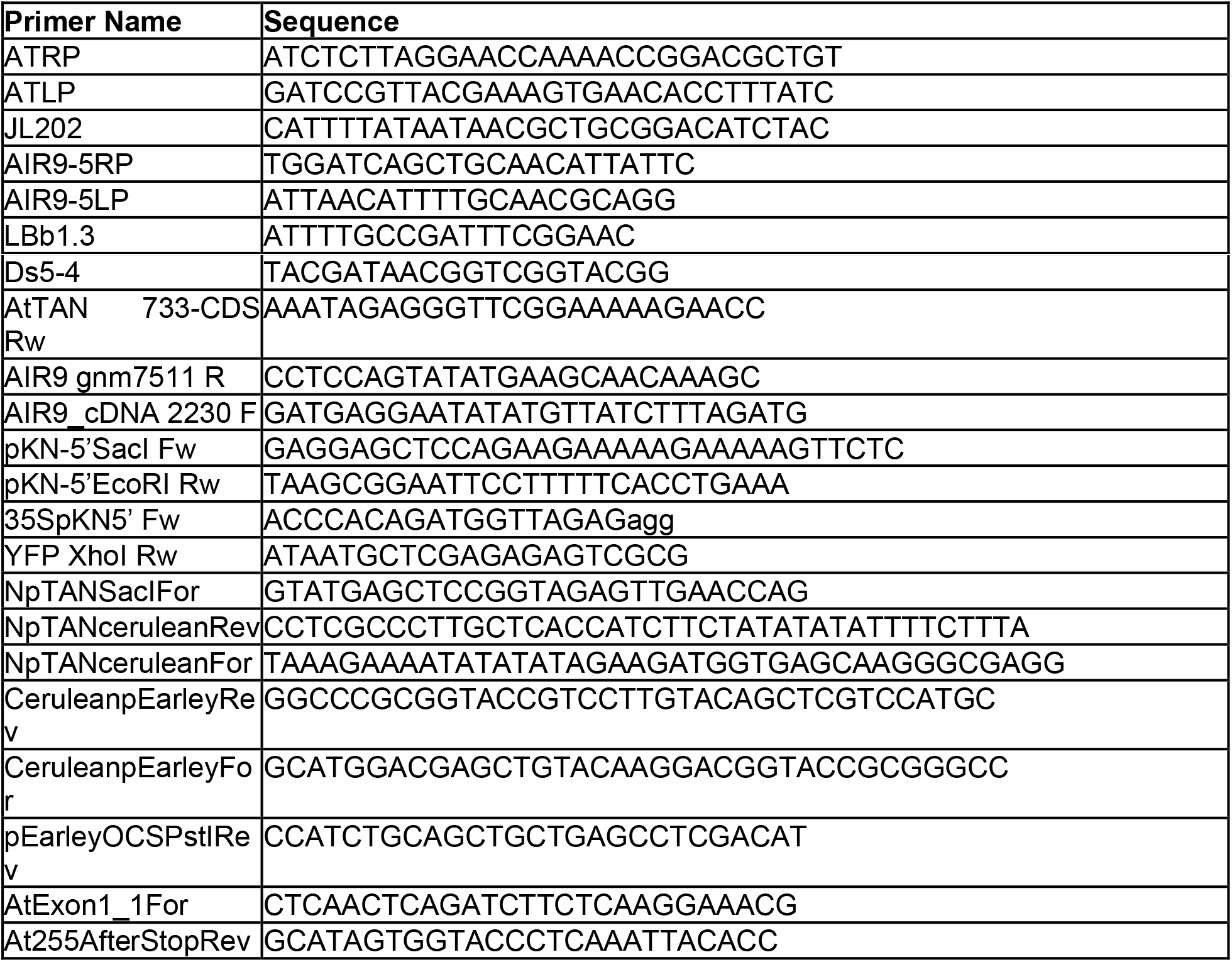
Primers used for cloning and genotyping.

